# Methionine coordinates a hierarchically organized anabolic program enabling proliferation

**DOI:** 10.1101/249367

**Authors:** Adhish S. Walvekar, Rajalakshmi Srinivasan, Ritu Gupta, Sunil Laxman

## Abstract

Methionine availability during overall amino acid limitation metabolically reprograms cells to support proliferation, the underlying basis for which remains unclear. Here, we construct the organization of this methionine mediated anabolic program, using yeast. Combining comparative transcriptome analysis, biochemical and metabolic flux based approaches, we discover that methionine rewires overall metabolic outputs by increasing the activity of three key regulatory nodes. These are: the pentose phosphate pathway coupled with reductive biosynthesis, and overall transamination capacity, including the synthesis of glutamate/glutamine. These provides the cofactors or substrates that enhance subsequent rate-limiting reactions in the synthesis of costly amino acids, and nucleotides, which are also induced in a methionine dependent manner. This thereby results in a biochemical cascade establishing an overall anabolic program. For this methionine mediated anabolic program leading to proliferation, cells co-opt a “starvation stress response” regulator, Gcn4p. Collectively, our data suggest a hierarchical metabolic framework explaining how methionine mediates an anabolic switch.

## Introduction

Cell growth is expensive, and is therefore tightly co-ordinated with the intrinsic cellular metabolic state. In general, the enormous metabolic costs incurred during growth and proliferation come from two well-studied phenomena. First, to successfully complete division, a cell makes substantial metabolic investments, in order to replicate its genome, and synthesize building blocks like amino acids, lipids, nucleotides, and other macromolecules^1^. Second, the process of protein synthesis required for growth itself consumes large amounts of energy^2,3^, as the translational output of cells increases^4^. Understanding such changes in cellular metabolic state coupled to global biosynthetic outputs, in the context of commitments to cell growth and proliferation are now the focus of several studies^5–9^. For example, one context where there is intense interest in understanding metabolic alterations enabling growth is in cancers, where phenomena ranging from the Warburg effect^10,11^, to the identification of the biosynthetic and metabolic requirements for cell growth^11–13^ are studied. Yet, given the overall complexity of metabolic rewiring, understanding how specific, “sentinel” metabolites can function directly as growth signals, and identifying the core, necessary steps by which such metabolites can reprogram cells to an anabolic state, has been challenging.

However, simple, tractable cellular models can be used to dissect and deconvolute such complex phenomena. Studies using *Saccharomyces cerevisiae* have been particularly instrumental in identifying dedicated, conserved strategies utilized by eukaryotic cells to integrate metabolic state with growth^5,6,8,9,14–19^. In such reductive studies using yeast, preferred carbon or nitrogen sources are typically limited (thereby slowing down overall growth). Subsequently, specific factors are reintroduced individually or in combination. This thereby reconstitutes minimal components required for reprogramming cells to an anabolic state, and allows the precise identification of necessary components, or dissection of mechanistic events. Such approaches have discovered novel nutrient sensing systems, and mechanisms by which growth outputs are controlled by the build-up and utilization of specific metabolites^8,9,25–27,15,17,18,20–24^.

Interestingly, some amino acids directly function as anabolic signals, potently activating growth pathways independent of their roles as nitrogen or carbon sources. For example, leucine and glutamine activate the TOR pathway directly^28,29^. In this context, studies from diverse organisms, observed over many decades suggest that methionine is a strong growth signal, or “growth metabolite”^25,30–36^. The most direct evidence for methionine as a growth signal come from recent studies in yeast. When *S. cerevisiae* are shifted from complex, amino acid replete medium with lactate as the carbon source, to a minimal medium with the same carbon source, the addition of methionine alone (likely through its metabolite S-adenosylmethionine (SAM)), strongly promotes growth and proliferation^17,25,26,37,38^. Thus, even during otherwise overall nutrient limitation, methionine can induce proliferation. Despite these advances, two fundamental, related questions regarding methionine as a growth signal remain unanswered. First, what is the biochemical logic of the methionine mediated anabolic program (i.e. how does methionine result in an anabolic reprogramming)? Second, what is the mechanism by which methionine mediates this anabolic rewiring, even in overall amino acid limiting conditions? We address these related questions in this study.

Here, using a minimal, reconstitutive system in yeast, and a biochemical first-principles approach, we uncover how methionine uniquely rewires cells to an anabolic state, even in otherwise amino acid limited conditions. We find that methionine activates very specific metabolic nodes in order to mediate this anabolic reprogramming. When these nodes are coincidently activated, they further induce a cascade of dependent metabolic processes leading to the overall biosynthesis of “costly” amino acids and nucleotides. For appropriately executing this anabolic program and sustaining proliferation, cells co-opt Gcn4p, a mediator of a nutrient stress/survival response. Collectively, these results position methionine at the apex of an overall anabolic network, and provide an overarching, hierarchically organized metabolic logic to understand how methionine availability results in metabolic rewiring and controlling cellular metabolic state.

## Results

### Methionine mediates a transcriptional remodelling program inducing key anabolic nodes

When wild-type, prototrophic yeast cells are shifted from a complex, amino acid rich medium with lactate as the sole carbon source (RM), to a synthetic minimal medium containing nitrogen base and lactate (MM), they show a significant lag phase and slower growth. Supplementation with all 20 standard amino acids restores growth after this nutrient downshift^25,38^. Importantly, methionine supplementation alone substantially increases growth (Figure 1A), comparable to (in our hands) or better than adding all eighteen other non-sulfur amino acids (nonSAAs) together^25,38^. Collectively, even during otherwise overall amino acid limitation, methionine availability increases proliferation. Given that methionine is itself not a good “nutrient source” (i.e. a poor carbon or nitrogen source), and adding methionine alone cannot create a nutrient replete medium, we wondered if methionine might mediate a complete switch to an anabolic state. This is a clear metabolic supply problem that needs to be solved. Thus, we reasoned that dissecting the methionine-mediated overall transcriptional response might provide insight into the logic of the anabolic program mediated by methionine, and allow the elucidation of a core metabolic response that drives proliferation.

**Figure 1:**
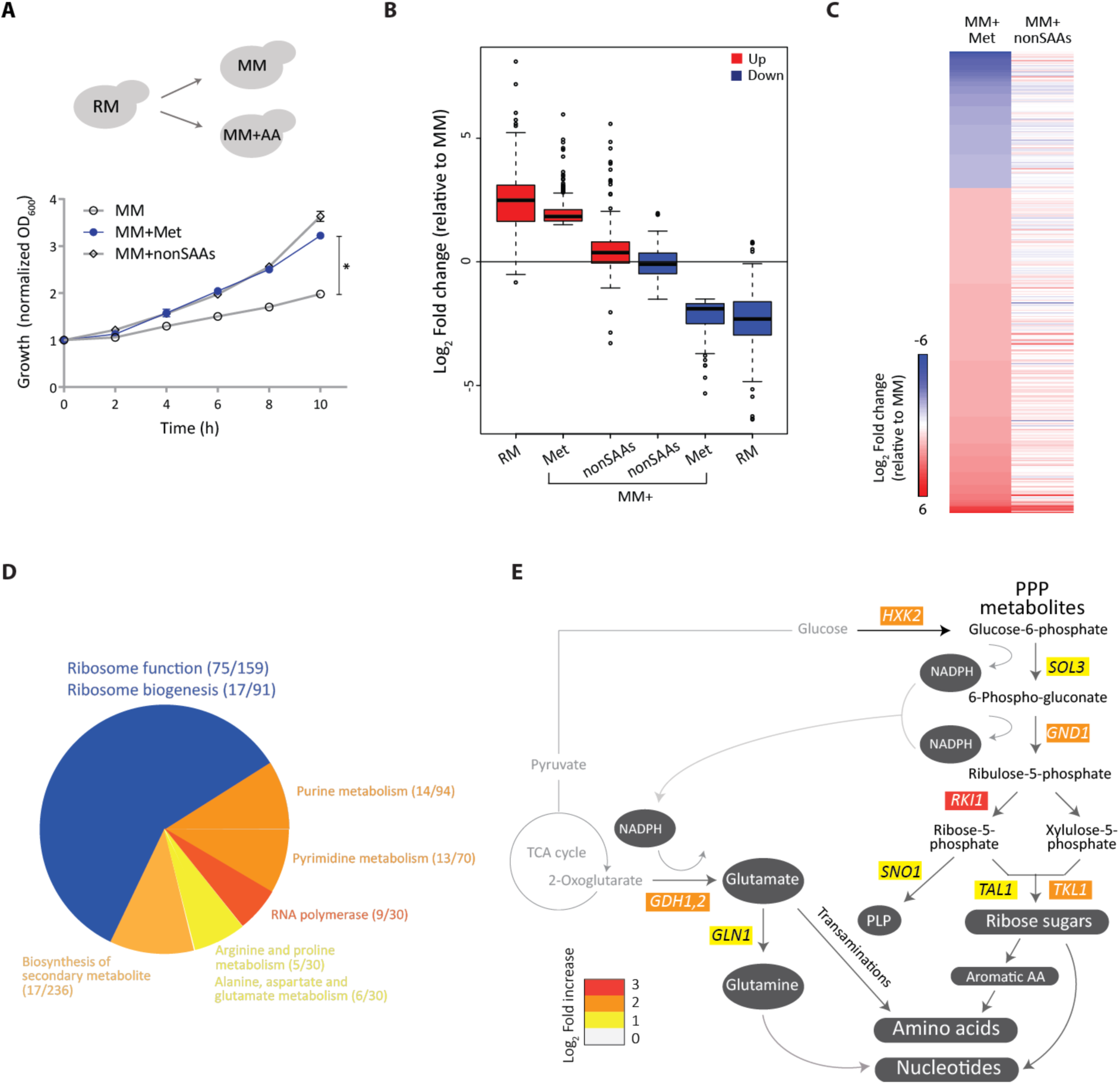
Methionine mediates a transcriptional remodelling program inducing key anabolic nodes. A) Methionine and cell proliferation during amino acid limitation. Shown are growth profiles of WT cells grown in rich medium (RM) and shifted to minimal medium (MM) with or without the indicated amino acid supplements (2 mM each; nonSAAs indicates all the non-sulfur amino acids except tyrosine). The growth profile with methionine is in blue. B) Global trends of gene expression in RM and methionine supplemented MM. The boxplot shows gene expression levels of transcripts in WT cells grown in MM plus methionine in comparison to the MM set, and compares the expression levels of these genes in the RM or MM plus nonSAAs sets. C) Effect of methionine on a transcriptional program in cells. The heat map shows differences in differentially expressed genes in cells grown in MM plus methionine compared to MM (left column), with cells grown in MM plus nonSAAs compared to MM (right column). D) Gene Ontology (GO) based analysis of the methionine-induced genes. The pie-chart depicts the processes grouped by GO analysis for the up-regulated transcripts between MM plus methionine and MM set. Numbers in the bracket indicate the number of genes from the query set/ total number of genes in the reference set for the given GO category. E) Manual regrouping of the methionine responsive genes into their relevant metabolic pathways, pertinent only to carbon metabolism and central metabolic processes. The pathway map includes individual genes in central carbon metabolism which are induced by methionine (indicating the fold-changes in gene expression). The arrows marked red indicate increased expression across the pathway. In all panels, data shows mean±SD. ^*^p < 0.05.

In this section we first address how methionine reprograms cells into an anabolic state, focusing on elucidating relatively early transcriptional events (before the overall proliferation is observed). We performed comprehensive RNA-seq analysis on distinct sample sets of wild-type cells- (i) RM grown, or cells shifted to (ii) MM for 2h, (iii) MM+Met for 2h (Met set), and (iv) MM+nonSAAs for 2h. Transcript reads from the biological replicates showed exceptional correlation across all conditions (Pearson correlation coefficient, R≥0.99) (Figure S1). Setting a stringent cut-off, we only considered differentially expressed genes with ≥ log_2_ 1.5 fold changes (i.e. ~2.8 fold change), and a p-value cut-off <10^−4^ for further analysis. We first compared global transcription trends in WT cells growing in RM, MM+Met or MM+nonSAAs to MM, with the focus being what happens when methionine is the sole variable. We examined overall global gene expression trends in these conditions (compared to MM), looking at the distribution of the most induced or downregulated genes (Figure 1B, Figure S2). Here, we first compared the global gene expression trends (broad trends of up- or down- regulated genes) exhibited by cells in MM+Met, to cells grown in RM or MM+nonSAAs, all relative to MM (i.e. we compared the expression profiles of the genes up/downregulated in MM+Met, to the same genes in RM or MM+nonSAAs, all baselined to these gene expression levels in MM) (Figure 1B). Notably, the MM+Met gene expression profile very closely resembled the signature of cells in RM, in contrast to the cells in MM+nonSAAs (which were nearly indistinguishable from MM) (Figure 1B, Figure S2 and S3). This suggests that methionine alone (compared to all other nonSAAs combined) is perceived by cells as a stronger anabolic cue than all non-sulfur amino acids combined, and is sufficient to switch cells into a transcriptional state resembling that of rapidly proliferating cells in RM (which is complex, amino acid rich media ideal for growth).

We next more closely examined the overall transcriptional response unique to methionine, by comparing transcriptomes of cells growing in MM vs MM+Met (the only variable being methionine). This comparison identified 372 genes, of which 262 genes were upregulated in the Met set (Figure 1C, Supplementary file E1). Using gene ontology (GO), these genes were grouped into related processes (Figure 1D, Figure S4A, Supplementary file E2). Given that there is an eventual growth increase in MM+Met, we expectedly observed a grouping suggesting a transcriptional induction of genes related to the core translational machinery. Additionally, GO also grouped multiple induced genes into “nucleotide metabolism”, i.e. under “purine/pyrimidine” or “nucleobase and nucleotide metabolism”, along with the biosynthesis of secondary metabolites (Figure 1D). All of this would be entirely expected for any cell in a “growth/proliferative” state, since proliferation relies on increased translation and replication. Unsurprisingly therefore, the GO grouping showed a signature of a cell in a “proliferative state”.

However, this form of GO based grouping does not resolve the metabolic supply problem highlighted earlier. This grouping does not address the underlying metabolic logic or hierarchy of the methionine mediated anabolic response and how it might have been achieved, but rather reveals the end-point readouts for growth. We speculated that this is due to a limitation of using GO based analysis, which builds groups by looking for enriched pathway terms relying on multiple genes within a pathway to be overrepresented. Contrastingly, for metabolic changes, entire metabolic pathways need not be regulated. This is because most metabolic regulation happens by controlling only key nodes or “rate-limiting steps” in metabolism^1^. We therefore manually rebuilt connections and groupings using biochemical first principles, in order to attempt to put together a logical hierarchy of metabolic responses being set-up. In this reconstruction, we particularly emphasized: (i) whether the protein encoded by the gene regulated a “bottleneck” or rate-limiting metabolic step, and (ii) whether this biochemical step, and its subsequent product, was critical for multiple other biosynthetic reactions. Contrastingly, we did not worry about how many genes in a pathway are induced (i.e. GO pathway enrichment). Our reasoning was that these bottleneck nodes will not necessarily be picked up in GO enrichments, particularly if they are solitary genes, and therefore entire metabolic pathways were not transcriptionally upregulated. However, these bottleneck genes may in fact be central to understanding the metabolic state switch.

Through this biochemical first-principles based reconstruction, we identified and compartmentalized the metabolic response regulated by methionine into a group of key biochemical reaction nodes, as described. Only a few genes involved in classical “central carbon/carbohydrate metabolism” were upregulated in the presence of methionine, and strikingly none of them group to glycolysis, the TCA cycle or gluconeogenesis (Figure 1E). However, the genes encoding three key enzymes of the pentose phosphate pathway (PPP) (*GND1, RKI1* and *TKL1*), which regulate four steps in the PPP, were induced in the presence of methionine. Furthermore, two other genes (*SOL3* and *TAL1*), which control two other steps in the PPP, were also induced by methionine (at just below the log_2_ 1.5 fold (~2.8 fold) arbitrary cut-off limit we had set) (Figure 1E). Gnd1p catalyzes the last step in the oxidative arm of the PPP, generating NADPH and producing ribulose-5-phosphate. Most of the genes of the non-oxidative arm of the PPP, which make ribose-5-phosphate and other critical intermediates, were also upregulated. Additionally, *HXK2*, encoding a hexokinase was upregulated when methionine is present. While this is not even classified under the PPP by GO grouping, this enzyme produces glucose-6-phosphate, which is the substrate for the first, rate-limited step of the PPP, and so we included it under the PPP in our grouping (Figure 1E). Thus, only the PPP arm of carbon metabolism was transcriptionally induced by methionine. Second, we noted that key regulator resulting in the formation of pyridoxal-5-phosphate or PLP (encoded by *SNO1*), was induced by methionine (Figure 1E). PLP is a central cofactor, required for all transamination reactions^1^, but notably does not get a GO grouping because it does not fall in a large pathway/group. Third, transcripts of Gdh1p, which regulates the key nitrogen assimilation reaction resulting in the formation of glutamate (and *GLN1*, which is further required to make glutamine), was highly induced in methionine (Figure 1E). This reaction requires NADPH (which is itself produced in the PPP), and importantly is also critical for the subsequent formation of all other amino acids, and nucleotides (Figure 1E). Thus, this grouping suggested that methionine induced a PPP-GDH-PLP metabolic node. We hypothesized that this PPP-GDH-PLP node could be central for all the subsequent, downstream anabolic outputs.

### Methionine sets-up a hierarchical metabolic response leading to anabolism

We therefore inspected our transcriptome data for the other genes upregulated by methionine, which could be grouped broadly under “amino acid biosynthesis”, “nucleotide synthesis”, and “oxidoreduction/transamination” categories, particularly focusing on the substrates or co-factors required for their function. Notably, only a few genes in each of these large, multi-step, multi-enzyme pathways were induced. However, when organized by their biochemical requirements, essentially all enzymes encoded by this set of methionine-upregulated genes utilized either a PPP intermediate/product, and/or NADPH, and/or glutamate/glutamine, or combinations of all of these, i.e. products coming from the PPP- GDH-PLP nodes (Figure 2A). We next more closely examined the steps in the respective biosynthetic pathways that these genes regulated (coming from Figure 2A). For this, we further categorized all steps in the amino acid biosynthesis pathway as either the rate-limiting/initiation step and/or final step in the production of that amino acid, and organized them based on the use of costly and complex precursors or cofactors (Figure 2B). Strikingly, we observed that the methionine induced genes (which are few in number) in these pathways regulated only the most critical, rate-limiting or costly steps in amino acid biosynthesis (p-value of 3.8e^-09^, Fisher’s exact test), but had little or no significant role in regulating the multiple other genes in the pathway that encode enzymes for inexpensive steps (Figure 2B). Finally, we note that these methionine-induced genes in the “amino acid biosynthesis” bin do not just broadly represent all amino acid biosynthesis, but synthesize what are viewed as the costliest amino acids to synthesize^39^, namely the aromatic amino acids, the branched-chain amino acids, and lysine, which is highly overrepresented in ribosomal and core translational machinery proteins^26^.

**Figure 2:**
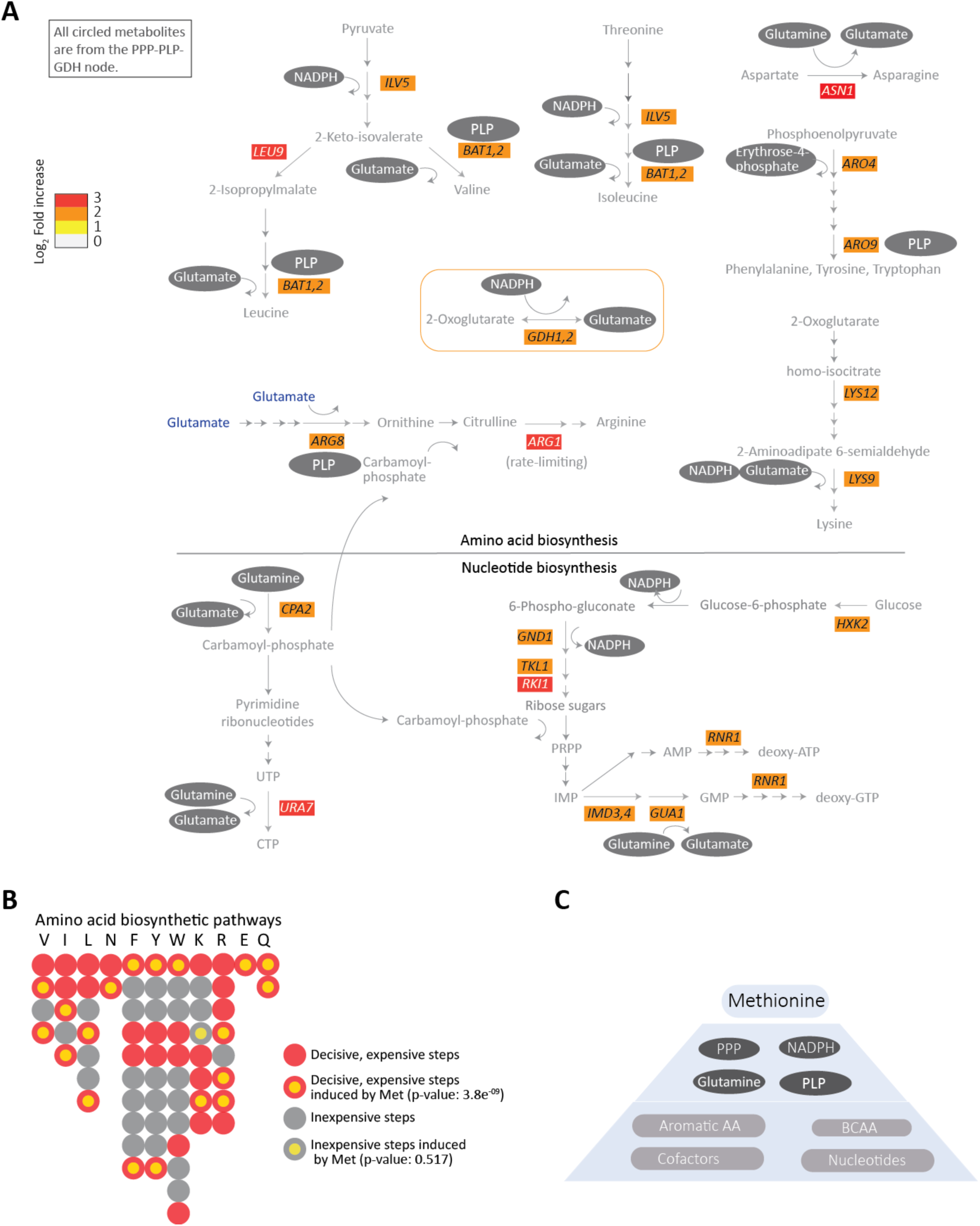
Methionine sets-up a hierarchical metabolic response leading to anabolism. A) Regrouping of the methionine induced genes, focusing on those directly involved in amino acid and nucleotide metabolism. The schematic shows the methionine-responsive genes in various amino acid and nucleotide biosynthesis pathways, along with their fold-changes in gene expression. The requirement of PPP metabolites, NADPH, PLP or glutamate/glutamine (see Figure 1E) for the individual steps is mapped on to the schematic. The arrows marked red are the steps induced in the presence of methionine. Note that all gene products induced by methionine in these pathways use PPP intermediates, NADPH, PLP and/or glutamine/glutamate (indicated within grey ovals) in their biochemical reactions. B) A bird’s eye-view of the amino acid biosynthesis steps regulated by methionine, with the metabolic costs associated with each step indicated. Each bead (or filled circle) represents a step in the pathway (prepared according to the individual amino acid pathways shown at https://pathway.yeastgenome.org/.) A step is considered expensive (marked red) when it is either the entry or the final or involves ATP utilization or involves reduction. All the rest of the steps are considered inexpensive (marked grey). Methionine induced steps are shown with a yellow fill at the centre of the circle, for the given step. The p-value for methionine dependence of genes encoding the critical, rate-limiting or costly steps in amino acid= 3.8e^-09^, and for the other nodes, it is non-significant (Fisher’s exact test). C) A hierarchical organization of the methionine mediated anabolic remodelling. Methionine induces expression of genes in the PPP-GDH-PLP node, which provides precursors for the key steps in the biosynthesis of all other amino acids and nucleotides, and these steps are also directly induced by methionine.

Similarly, nucleotide biosynthesis involves very elaborate, multi-step, multi-enzyme pathways. Using a similar logic to group pathways, we find that the methionine dependent, upregulated genes again encoded enzymes controlling very specific, limiting steps in nucleotide synthesis (Figure 2A). Furthermore, these regulated steps all utilize glutamate/glutamine and/or NADPH, as well as pentose sugars from the PPP, i.e. the PPP- GDH-PLP nodes (Figure 2A). Notably, *RNR1*, which encodes the key enzyme in converting ribonucleotides to deoxyribonucleotides (and hence the critical hub for DNA synthesis), is strongly upregulated upon methionine addition (Figure 2A), while most other steps (which are not rate limiting) are not regulated by methionine. Separately, as a control, we expanded this analysis and compared the methionine response to minimal medium supplemented with all other nonSAAs, and here the overall metabolic grouping or organization (for methionine induced genes) remained unchanged (Figure S2), with nonSAAs resembling MM. Finally, in MM+nonSAAs, the few highly induced genes (compared to RM) function in methionine (and sulfur-amino acid) related biosynthesis or salvage (Figure S4B, Supplementary file E1), and not additional reactions. This further substantiates our overall observations for the role of methionine as an “anabolic signal”.

Collectively, this comparative transcriptome based analysis, focusing on the methionine induced metabolic program and carried out using a biochemical first principles approach, suggests not just a general anabolic remodelling due to methionine, but a hierarchical metabolic organization induced by methionine (Figure 2C). In this putative hierarchical organization, methionine induces genes regulating the PPP, key transamination reactions, and the synthesis of glutamine/glutamate (the PPP-GDH-PLP node). These three processes directly allow critical steps in synthesis of the costliest amino acids and nucleotides. The key, limiting steps in these subsequent synthesis reactions are themselves induced by methionine, collectively setting up a structured anabolic program (Figure 2C). These data thus uniquely position methionine as an anabolic cue.

### The core metabolic response induced by methionine is regulated by *GCN4*

How might methionine mediate this very specific transcriptional response to induce these metabolic nodes and genes? We reasoned that there must be a methionine dependent activation of a transcriptional regulator(s), which can specifically induce these metabolic genes, including amino acid and nucleotide biosynthetic genes. Further, the methionine effect was strongest in conditions of overall amino acid limitation. While there is currently no known methionine dependent transcriptional regulator that can control these metabolic nodes, there is in fact a well-known master-regulator of amino acid biosynthesis. The conserved transcription factor Gcn4p (Atf4 in mammals) is a transcriptional activator, primarily controlling the amino acid biosynthetic genes during amino acid starvations^40^. Although the activity of Gcn4p has been mainly studied during starvation as a “stress response” regulator, and not in contexts involving increased proliferation, we wondered if a possible connection between methionine and Gcn4p might exist. We therefore first monitored the amounts of endogenous Gcn4p (chromosomally tagged with a C-terminal HA epitope) after a shift to MM, with and without supplementation of different amino acids including methionine. Surprisingly, Gcn4p amounts increased substantially specifically upon methionine supplementation alone (when other amino acids were not supplemented), compared to either MM, or MM supplemented with all 18 other nonSAAs (Figure 3A). This observation was independently confirmed using immunofluorescence based experiments (Figure S5A). We therefore asked whether Gcn4p was necessary for the increased growth upon methionine supplementation. Notably, the *gcn4Δ* cells did not show any increased growth in methionine supplemented medium, but instead grew comparably to WT cells in MM (Figure 3B). As controls, in all other conditions (lacking methionine, or in RM) the growth of *gcn4Δ* cells was indistinguishable from the WT cells (Figure S5B). Collectively, these data suggest that Gcn4p is necessary for the methionine-mediated growth in otherwise amino acid poor conditions. We therefore hypothesized that the methionine dependent transcriptional response, particularly those related to metabolism, might be mediated by Gcn4p. To address the role for Gcn4p in this anabolic response, we next carried out a comparison of transcriptomes of *gcn4Δ* cells grown in RM, MM, MM+Met and MM+nonSAAs with wild-type cells grown in the respective conditions.

**Figure 3:**
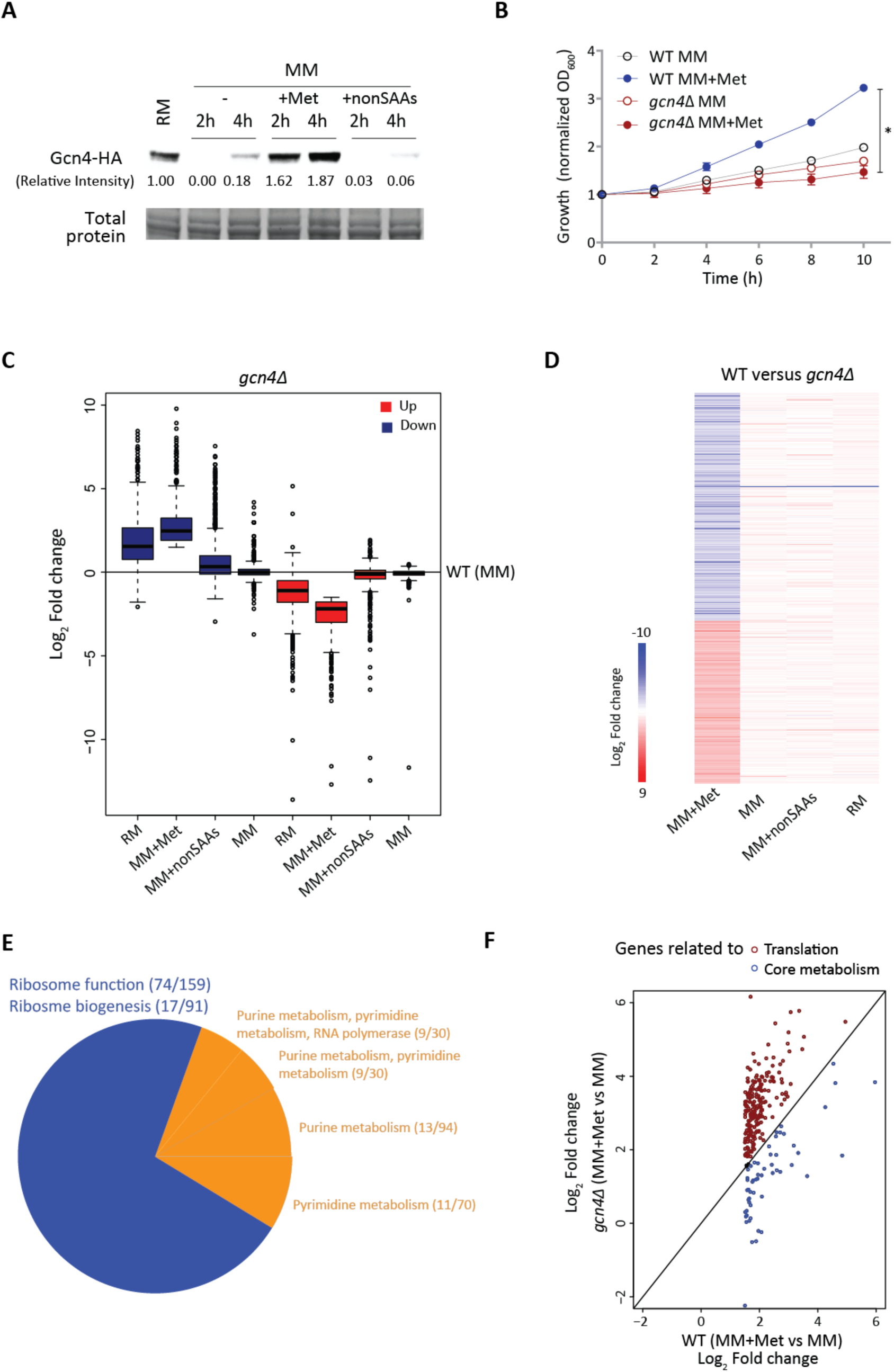
The core metabolic response induced by methionine is regulated by *GCN4*. A) Gcn4p is induced by methionine. Gcn4p amounts were detected by Western blotting of WT cells (expressing Gcn4p with an HA epitope, tagged at the endogenous locus) shifted from RM to MM, or MM supplemented with the indicated combinations of amino acids. A representative blot is shown. B) *GCN4* is necessary for methionine-mediated increased growth. WT and *gcn4Δ* cells were shifted from RM to MM with or without methionine supplementation and growth was monitored. Also see Figure S5B. C) Trends of gene expression in RM and methionine supplemented MM in *gcn4Δ* cells. Gene expression levels of transcripts in *gcn4Δ* cells grown in RM or shifted to MM or MM plus methionine or MM plus nonSAAs were compared to WT MM set. D) Global transcriptional response in the absence of *GCN4*. The heat map shows differentially expressed genes (log_2_ 1.5-fold change; p < 10–^4^) between WT and *gcn4Δ* cells in the respective growth conditions. E) GO based analysis of the methionine-responsive genes in *gcn4Δ* cells. A pie chart showing the processes grouped by GO analysis for the up-regulated transcripts between MM plus methionine and MM set in *gcn4Δ* background. Numbers in the bracket indicate the number of genes from the query set/ total number of genes in the reference set for the given GO category. F) The metabolic program is *GCN4* dependent. Categorization of the *GCN4* dependent transcripts in the presence of methionine, as related to metabolism, or translation. The expression level of the methionine-responsive transcripts related to metabolism and translation in WT set (MM plus methionine versus MM) was compared with the *gcn4Δ* background. The genes related to the metabolic steps described in Figures 1 and 2 are marked with blue circles, while genes related to ribosome biogenesis and function are marked with red circles. In all panels, data shows mean±SD. ^*^p < 0.05.

We first examined global trends of gene expression (similar to those in Figure 1B) in the absence of Gcn4p, and compared those to the WT set. Here, the baseline again was gene expression in WT cells in MM. To our surprise, the overall global gene expression trends in the Met set (in *gcn4Δ* cells) even more strongly resembled the RM set than WT cells (in methionine, from Figure 1B) (Figure 3C). The transcriptional response in all the other conditions (RM or nonSAA) was almost unaffected in *gcn4Δ* cells compared to WT cells (Figure 3C). This paradoxically suggested that in the absence of *GCN4*, methionine invokes an even stronger transcriptional response, with global trends seemingly resembling a strong “growth state” (Figure 3C). We also more closely compared transcriptomes of WT cells with *gcn4Δ* cells, under the same combination of conditions used earlier, and analyzed our data with the stringent filters used in the previous section. The overall changes in transcriptomes of WT vs *gcn4Δ* cells are shown (Figure 3D, Figure S6, Supplementary file E1). In all conditions except methionine, WT and *gcn4Δ* cells showed very similar gene expression profiles (Figure 3D), suggesting a unique role for Gcn4p in the presence of methionine. To better understand what component of the methionine response was directly induced in a Gcn4p dependent manner, we organized the genes induced in methionine (in the *gcn4Δ* cells, compared to WT cells in MM) by function. When grouped using GO, we found that a very large number of genes (~200) induced in the presence of methionine, grouped into the general groups of “ribosome/translation”, and “nucleotide synthesis” (Figure 3E, Supplementary file E2). Surprisingly, this representation was even more striking than that seen in WT cells (Figure 1D), suggesting a strong “growth signature” in the presence of methionine even in cells lacking Gcn4p.

All these data suggested the perplexing observation that cells lacking Gcn4p showed a very strong transcriptional response in MM+Met, and that the signature of this transcriptional response largely remained similar to, but stronger than WT cells in MM+Met. Note however that (as shown earlier in Figure 3B) Gcn4p was essential for the growth induction due to methionine. These data therefore paradoxically suggested that while the overall “growth signature” response due to methionine remained, and was being further amplified in the absence of Gcn4p, only a small subset of Gcn4p regulated genes may be pivotal for the growth outcome. Do recall that the increased growth in methionine was entirely Gcn4p dependent. We therefore more carefully analyzed the classes of transcripts induced in WT and *gcn4Δ* cells by methionine (MM+Met set), and separated our functional groupings (based on GO) into two broad bins. One bin represented all genes related to ribosome function and translation (as seen in Figure 1D), while the other bin separated out the core metabolic genes similar to those classified in Figures 1 and 2 (Figure 3E). Strikingly, transcripts of every gene that mapped to ribosome/translation function (from Figure 1D) was higher in *gcn4Δ* cells compared to WT cells in the presence of methionine (Figure 3E). Contrastingly, transcript amounts of every gene related to amino acid biosynthesis, the PPP and nucleotide metabolism (and related metabolism) was significantly lower in the *gcn4Δ* cells than in WT cells (Figure 3E). Thus, this reorganization revealed that in the presence of methionine, while the induction of the translation machinery genes (which is the “growth signature”) was not Gcn4p dependent, the entire methionine induced core anabolic program hinged upon Gcn4p.

### Methionine induced enzymes in amino acid and nucleotide biosynthesis are Gcn4p dependent

We therefore more systematically examined the methionine and Gcn4p dependent transcription of genes that functionally regulated the PPP-GDH-PLP node, or specific steps in amino acid biosynthesis, which constitute the metabolic hierarchy we have described (Figure 4A). Strikingly, the genes from the PPP, and transamination reactions were strongly downregulated in *gcn4Δ* cells compared to WT cells in methionine (Figure 4A). However, the transcript of the Gdh1 enzyme (required for glutamate synthesis) was only methionine, but not Gcn4p dependent (Figure 4A). Next, the transcripts of methionine induced genes encoding the key, rate-limiting steps in multiple branched-chain or aromatic amino acid, and lysine and arginine biosynthesis were also Gcn4p dependent. Finally, while most nucleotide biosynthesis genes were not Gcn4p dependent, the entire methionine induced RNR complex (which is critical for the NTP to dNTP conversion, required for DNA synthesis) was strongly Gcn4p dependent (Figure 4A). Collectively, these data suggest that the core anabolic program induced by methionine is Gcn4p dependent.

**Figure 4:**
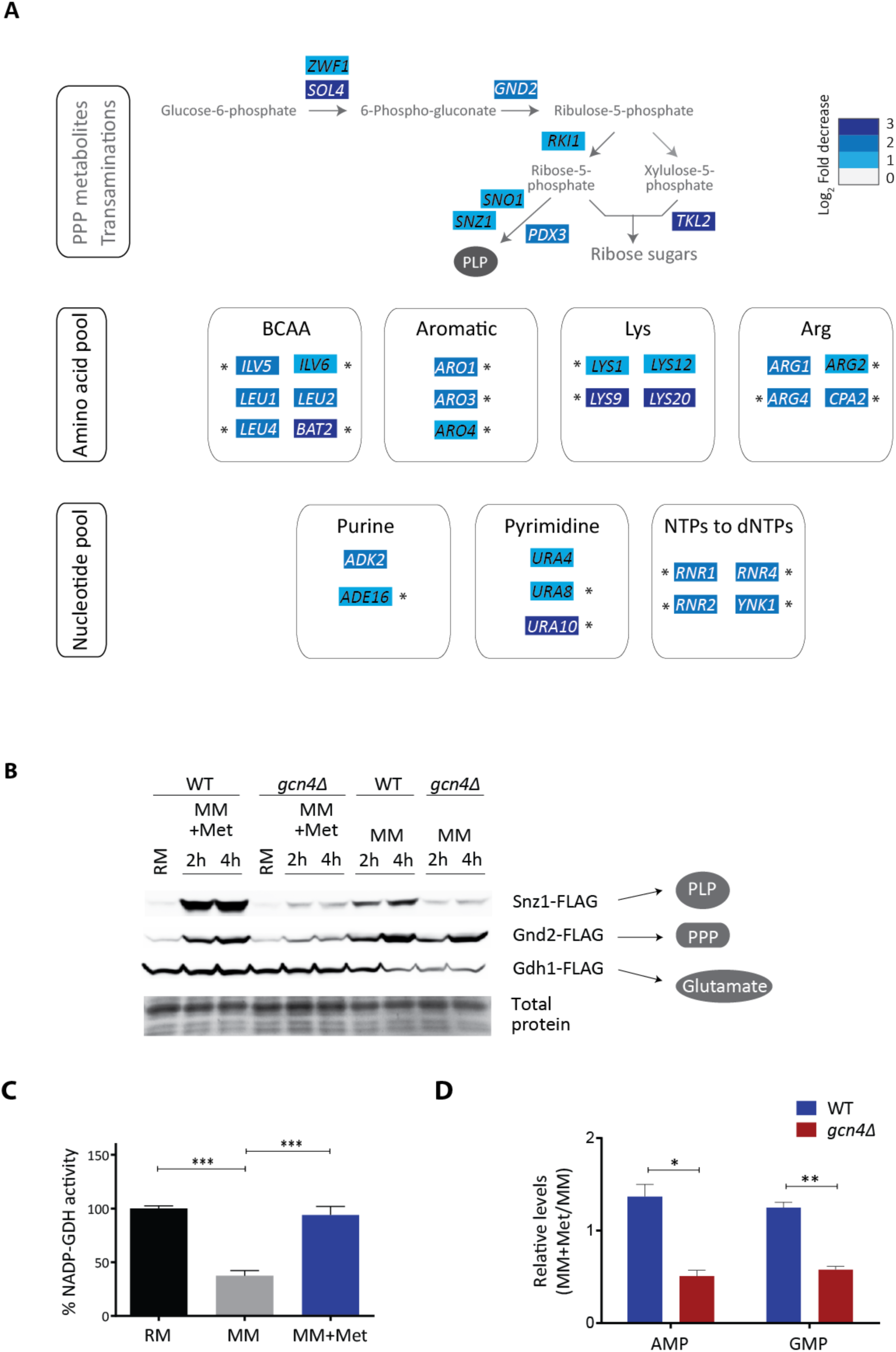
Methionine induced enzymes in amino acid and nucleotide biosynthesis are Gcn4p dependent. A) *GCN4* regulates the metabolic program due to methionine. Regrouping of the *GCN4*-dependent genes based on the PPP-PLP-GDH dependent metabolic nodes. The schematic shows the *GCN4*-dependent genes (comparison of MM plus methionine set between WT and *gcn4Δ*) in the PPP, amino acid and nucleotide biosynthesis pathways, along with fold-changes in gene expression. The arrows marked blue in the PPP pathway are the steps down- regulated in *gcn4Δ* cells. The rate-limiting steps in the pathway are marked by asterisk. B) Snz1p, Gnd2p and Gdh1p amounts in WT or *gcn4Δ* cells, with methionine as a variable. WT and *gcn4Δ* cells expressing FLAG-tagged Snz1p or Gnd2p or Gdh1p were shifted from RM to MM or MM plus methionine and amounts of these proteins were detected by Western blotting. A representative blot is shown in each case. C) NADP-dependent glutamate dehydrogenase activity with methionine as a variable. Crude extracts of WT cells grown in RM and shifted to MM or MM plus methionine were analysed for intracellular biosynthetic NADP-glutamate dehydrogenase activity. D) Relative nucleotide amounts in the presence of methionine in WT or *gcn4Δ* cells. WT and *gcn4Δ* cells grown in RM were shifted to MM (4h) with and without methionine, and the relative amounts of AMP and GMP from metabolite extracts of the respective samples were measured by LC-MS/MS. In all panels data indicate mean±SD. ^*^p < 0.05, ^**^p < 0.01, ^***^p < 0.001.

We next directly assayed the extent of (i) methionine controlling this anabolic rewiring, through the PPP-GDH-PLP node feeding subsequent anabolic reactions; and (ii), the extent of Gcn4p dependence for these steps. We first tested this at the biochemical level, comparing the enzyme amounts of three targets that represent the coupling of the PPP with other processes (the PPP-PLP-GDH node). These are Snz1p, Gnd2p and Gdh1p. Snz1p is required for pyridoxal phosphate (PLP) biosynthesis^41^, which is essential for all transamination reactions^1^. As illustrated earlier in Figure 1E, PLP biosynthesis itself also requires the PPP intermediate erythrose-4-phosphate as a substrate. Gnd2p is the key NADPH generating enzyme in the oxidative branch of the PPP. Gdh1p consumes NADPH and makes glutamate from 2-ketoglutarate. We measured amounts of these three proteins from WT and *gcn4Δ* cells growing in MM, and MM+methionine (Figure 4B). Notably, Snz1p and Gnd2p showed a strong induction that was both methionine dependent, and dependent on Gcn4p (Figure 4B, Figure S7). Gdh1p was strongly dependent upon methionine, but was not dependent on Gcn4p (Figure 4B). We also measured *in vitro* Gdh1p activity (NADPH-GDH activity) in lysates from cells growing in MM or with methionine, and found that overall Gdh1p activity was higher in cells growing with methionine (Figure 4C). As seen in the Western blot analysis (Figure 4B), the *in vitro* Gdh1p activity was not dependent on Gcn4p (not shown). All these data strongly support the observation that methionine drives these key, coupled steps in biosynthesis, and that they are largely mediated by Gcn4p. Finally, an expected the final readout of this biochemical coupling should be changes in steady-state nucleotide amounts, with a Gcn4p dependence in methionine replete conditions. Comparing relative amounts of nucleotides in wild-type and *gcn4Δ* cells, we noted a decrease in the nucleotide amounts in *gcn4Δ* cells in the presence of methionine (Figure 4D). Collectively, these direct biochemical read-outs support our proposed paradigm of a coupled induction of the PPP, and key transamination reactions by methionine, leading to increased amino acid and nucleotide synthesis.

### Methionine increases amino acid biosynthesis *in vivo*

Substantiating these findings using steady-state metabolite measurements alone is insufficient (and often misleading), since any steady-state metabolite measurement (as in Figure 4D) cannot directly distinguish synthesis from consumption. Therefore, to directly address this possible hierarchical anabolic program, we resorted to a stable-isotope pulse labelling and an LC-MS/MS based approach to directly measure the new synthesis of amino acids. To WT or *gcn4Δ* cells in the respective medium with or without methionine, we pulsed ^15^N-labelled ammonium sulfate, and measured the ^15^N incorporation into amino acids (Figure 5A, Table S2), before an effective steady-state of labelled amino acid synthesis and consumption was reached. This permits the detection of newly synthesized amino acids, which will incorporate the ^15^N label. We observed that biosynthesis of all the aromatic amino acids, lysine, histidine, proline, arginine and asparagine is strongly dependent on methionine presence (Figure 5B). For technical reasons we could not measure label-incorporation into branched-chain amino acids. Notably, the label immediately (~20 min) percolated in asparagine and aromatic amino acid biosynthesis, and showed a very strong methionine dependence (Figure 5B). Asparagine, proline and phenylalanine biosynthesis were methionine dependent even in *gcn4Δ* cells, pointing towards possible *GCN4*-independent influences of methionine. For all the other amino acids measured, the biosynthesis was both methionine as well as *GCN4*-dependent (Figure 5B). These data directly indicate that methionine availability controls the key nodes around PPP-PLP-GDH axis, thereby generating the amino acid pool required for proliferation, and that this is largely regulated by Gcn4p.

**Figure 5:**
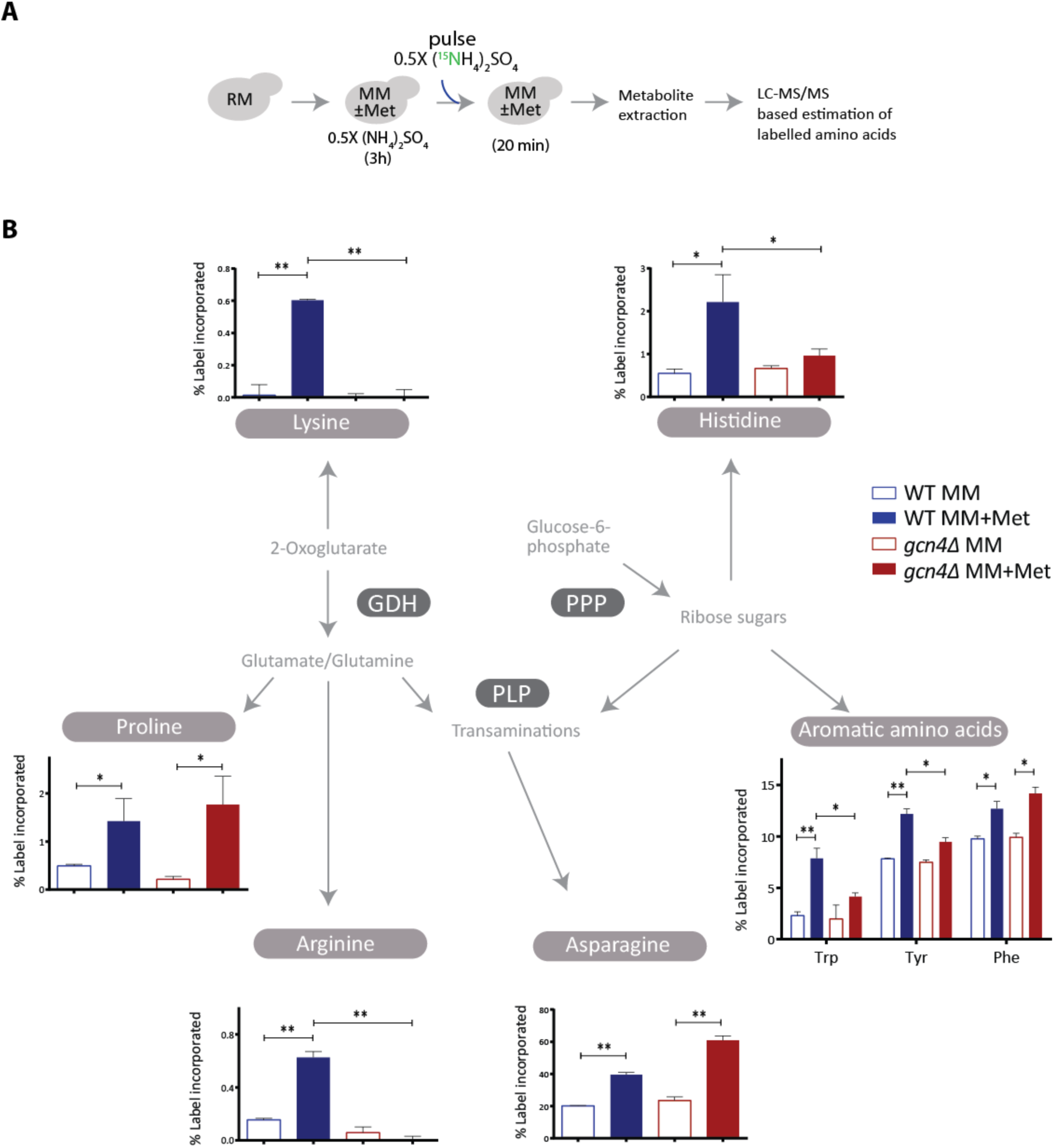
Methionine increases amino acid biosynthesis *in vivo*. A) A schematic showing the experimental design of ^15^N pulse-labelling experiment to measure amino acid biosynthetic flux. Cells were shifted to MM with and without methionine, maintained for 3h, and ^15^N-ammonium sulfate was pulsed into the medium, and the indicated, labelled metabolites were measured. Also see Table S2. B) Methionine increases amino acid biosynthesis in a Gcn4p dependent manner. ^15^N label incorporation into newly synthesized amino acids in WT and *gcn4Δ* cells was measured, as shown in the panel A. For all the labelled moieties, fractional abundance of the label was calculated. Also see Table S2 for mass spectrometry parameters. In all panels data indicate mean±SD. ^*^p < 0.05, ^**^p < 0.01.

### Methionine increases nucleotide biosynthesis *in vivo*

Given that the PPP and amino acid biosynthesis are directly regulated by methionine and Gcn4p, and the PPP metabolites and amino acids together couple to nucleotide synthesis, we predicted that collectively, perturbing this node should have a severe consequence on nucleotide biosynthesis. In principle, this also will reflect flux through the coupled steps of the PPP, glutamate/glutamine synthesis, and the use of intermediates from amino acid biosynthetic pathways for carbon and nitrogen assimilation into nucleotides (Figure 6A). The carbon skeleton of nucleotides comes from the PPP, the nitrogen base is directly derived from glutamine/glutamate and aspartate, and glutamate synthesis is itself coupled to NADPH (from the PPP) (Figure 6A). Nucleotide biosynthesis is also coupled to histidine and tryptophan synthesis. We therefore adopted a direct estimation of methionine and Gcn4p dependent increases in nucleotide synthesis (similar to the approach in Figure 5), predicting an increase in *de novo* nucleotide synthesis due to methionine, coming from the earlier amino acid precursors. To this end, using a stable-isotope based nitrogen or carbon pulse labelling approach, coupled to targeted LC-MS/MS based measurement of nucleotides, we separately measured the incorporation of the nitrogen and carbon label into nucleotides, as illustrated in Figure 6B and 6C. We observed a strong increase in ^15^N-labelled nucleotides upon the addition of methionine, in ~1 hour (Figure 6B, Table S2). Furthermore, this methionine-mediated incorporation of ^15^N-label in nucleotides was entirely Gcn4p dependent (Figure 6B and Figure S8).

**Figure 6:**
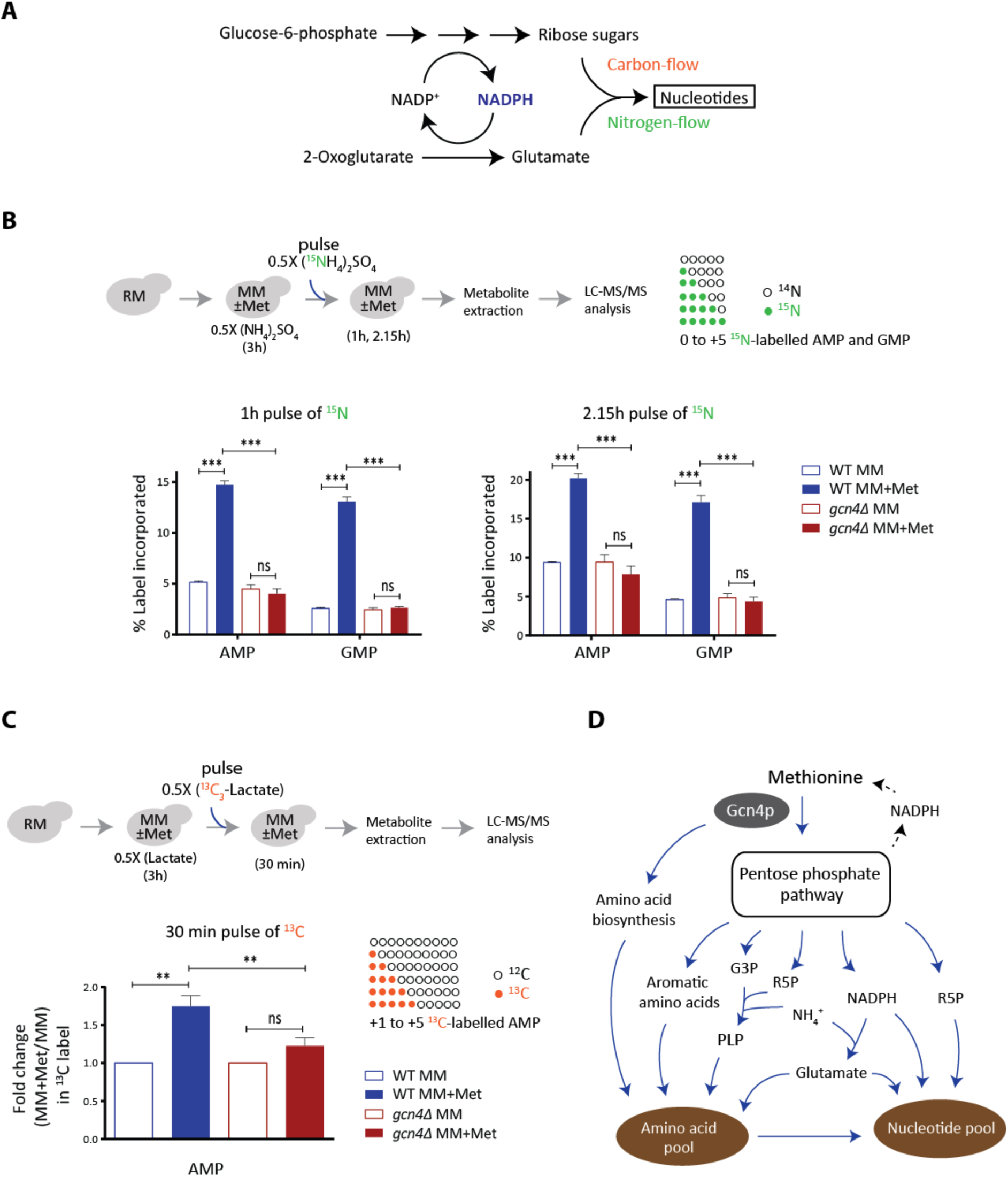
Methionine increases nucleotide biosynthesis *in vivo*. A) A schematic showing carbon and nitrogen inputs in nucleotide biosynthesis, and their coupling to the PPP/NADPH metabolism. B) Methionine increases nucleotide biosynthesis in a Gcn4p dependent manner. The WT and *gcn4Δ* cells treated and pulse-labelled with ^15^N ammonium sulfate as illustrated in the top panel. For all the labelled moieties, fractional increase of the incorporated label was calculated, to measure newly synthesized AMP and GMP. (also see Figure S8 for CMP and UMP). C) Methionine enhances carbon flux into AMP biosynthesis. An experimental set-up similar to the Panel B was employed, using ^13^C-lactate for carbon labelling. Label incorporation into nucleotides (from +1 to +5) was accounted for calculations. (note: GMP could not be estimated because of MS/MS signal interference from unknown compounds in the metabolite extract). D) A model illustrating how methionine triggers an anabolic program leading to cell proliferation. Methionine promotes the synthesis of PPP metabolites, PLP, NADPH and glutamate (up-regulated genes in the pathways are shown in blue), which directly feed into nitrogen metabolism. As a result, methionine activates biosynthesis of amino acids and nucleotides, allowing the cells to grow in amino acid limiting medium. In all panels data indicate mean±SD. ns: non-significant difference, ^**^p < 0.01, ^***^p < 0.001.

Monitoring carbon flux is extremely challenging in a non-fermentable carbon source like lactate (as compared to glucose), given the difficulties of following the labelled carbon molecules. Despite that, like the ^15^N-labeling experiments described above, a similar experimental design was adopted to measure the ^13^C-label incorporation into AMP (Figure 6C, Table S2). We observed a significant increase in ^13^C-labelled AMP upon the addition of methionine, and this methionine dependent incorporation of ^13^C-label in AMP was not observed in cells lacking Gcn4p (Figure 6C). Collectively, these data show a methionine and Gcn4p dependent increase in *de novo* synthesis of nucleotides, coupling carbon and nitrogen flux that is dependent on the PPP, and glutamate synthesis. Note that the overall kinetics of incorporation of label are entirely in line with the predicted hierarchy. Increased amino acid labels (shown in Figure 5) were seen in ~20 min post labelled ammonium sulfate addition, while the nucleotide label increase occurs in ~1 h, subsequent to the observed amino acid label increase. Thus, we directly demonstrate first the synthesis of new amino acids, and the subsequent synthesis of nucleotides, in a methionine and Gcn4p dependent manner.

## Discussion

In this study, we show how methionine drives cellular proliferation by rewiring cells to an anabolic state, even under otherwise amino acid limited, challenging conditions. We uncover a regulated, hierarchical activation of metabolic processes by methionine, which leads to overall anabolism. We also present a mechanism of how methionine mediates this anabolic program.

Starting with a global transcriptome analysis (Figure 1), we systematically build the underlying metabolic foundations of a methionine mediated anabolic state switch. Methionine mediates a global transcriptional remodelling in cells, thereby controlling the anabolic program (Figures 1 and 2). For understanding the core metabolic logic within the transcriptional response to methionine we adopted a biochemical first principles based approach, emphasizing control points at rate or resource limiting biochemical steps, instead of solely relying on standard GO based organization. The organizational metabolic logic that emerged was striking (Figure 1 and 2). First, methionine positively regulates the PPP (Figure 1). The PPP provides the pentose sugar backbones for nucleotides, along with reducing equivalents (NADPH), allowing reductive biosynthesis for a variety of anabolic molecules^1^. For amino acid and nucleotide synthesis, pyridoxal phosphate, which controls all transamination reactions, is also essential^42^, and methionine directly induced this node as well (Figure 1). Collectively, methionine strongly induced the PPP-GDH-PLP node (Figure 1). These three nodes can feed all the subsequent metabolic steps induced by methionine. Furthermore, in these subsequent anabolic nodes (the synthesis of amino acids and nucleotides), methionine only induces the expression of genes that control rate limiting or final steps (Figure 2). These biosynthetic nodes include those that synthesize what are considered the “costliest” amino acids, namely the aromatic amino acids and the branch chain amino acids (Figure 2). Notably, essentially every one of these (methionine regulated) steps use cofactors or intermediates from the PPP-GDH-PLP node (Figure 1 and 2). Thus, it appears that methionine sets up this striking metabolic hierarchy, as illustrated in a schematic in Figure 2C and 6D.

Interestingly, the methionine dependent growth and increased activity in most of these metabolic nodes, and thus the overall anabolic program, depends upon Gcn4p (Figure 3 and 4). Gcn4p is best understood as a regulator of amino acid biosynthesis during starvation^43^. Indeed, many of the *GCN4* targets picked up in our study compare well with the landmark study of *GCN4* targets^43^ (see Figure S9 and Table S3). However, this role of Gcn4p in the presence of methionine in synchronously controlling these key hubs of the PPP, transamination reaction, glutamate biosynthesis, coupled with rate-regulating steps in costly amino acid and nucleotide biosynthesis has not been previously appreciated, and we think this is central to the anabolic program resulting in increased cell proliferation. This is in contrast to the well-studied role for Gcn4p for survival during starvation, allowing a restoration of amino acid levels. Also interestingly, Gcn4p appears to regulate only the core metabolic program induced by methionine, and not the induction of the translation machinery (as seen in Figure 3). This induction of translation due to methionine might be through other mechanisms, including activation of the TOR pathway^25,44^, and there seems to be a separation of the methionine sensing machinery from the actual effector of the anabolic program (Gcn4p). Finally, a combination of rigorous biochemical, and metabolic flux based analysis using stable-isotopes directly demonstrate this hierarchical coupling of the PPP, NADPH utilization and transamination reactions (in both nitrogen assimilation and carbon assimilation) first towards the increased synthesis of aromatic and branch-chain amino acids, and next towards nucleotides, in a methionine and Gcn4p dependent manner (Figures 5 and 6). Collectively, our data permits the construction of an overall pyrimidical hierarchy of metabolic events, mediated by methionine, to set up an anabolic program (model in Figure 2C). This suggests more general organizational principles by which cells can specifically rewire metabolism.

The central role of the PPP in anabolism is now text-book knowledge^1^. Yet, a better appreciation of the importance of the PPP in mediating an anabolic rewiring is now emerging due to the association of the PPP to cancer metabolism^45,46^. Many anabolic transformations require contributions from the PPP, however, the metabolic cues regulating the PPP (and coupling to other processes) are not often obvious. Additionally, these studies ignore or underplay coincident but necessary metabolic events for proliferation. Our study directly addresses how methionine (likely through its downstream metabolite SAM), acts as an anabolic signal for cells, through setting up of the metabolic hierarchy explained earlier, with the co-incident PLP and glutamine nodes being critically important. This striking role of methionine regulating an anabolic program seems analogous to another central metabolite, acetyl-CoA, which is better known to determine cellular decisions towards growth^15,47–50^. Interesting correlations can be made from our observations to known roles of methionine in cancer cell metabolism, and metazoan growth. The earliest observations of methionine as important for proliferation in some cancers dates back to the 1950s^33–35^, and several types of cancer cells are addicted to methionine^33,51–58^. Other, distinct studies show that *Drosophila* fed on methionine rich diets exhibit rapid growth, high fecundity, and shorter lifespans^30–32^, all hallmarks of what a “proliferative” metabolite will do. Studies from yeast show how methionine inhibits autophagy, or regulates the TORC1 to boost growth^25,26,38^. One of the earliest known cell cycle entry check-points found in yeast links to methionine^59^. Upon sulfate (and thereby methionine) starvation, yeast cells arrest their growth to promote survivability^20^, and transform their proteome to preferentially express proteins containing fewer cysteine/methionine residues to save sulfur^60^. There are other, less appreciated observations connecting methionine metabolism and the PPP. Yeast cells lacking *ZWF1* (encoding glucose-6-phosphate dehydrogenase, the first enzyme in the PPP) exhibit methionine auxotrophy^61^, and methionine supplementation also increases the oxidative stress tolerance of *zwf1Δ*^62^. Despite these studies highlighting a critical role of methionine, such a hierarchical logic explaining the organizational principles of the anabolic program mediated by methionine, and the mechanisms by which this is mediated, has thus far been elusive. Our study provides this.

Our use of a “less-preferred” carbon source, lactate, has helped reveal regulatory phenomena otherwise hidden in glucose and amino acid rich laboratory conditions, where a surfeit of costly metabolic resources (for example, unlimited PPP intermediates) are present. Tangentially, several recent reports emphasize the importance of lactate as a carbon source in rapidly proliferating cells^63–65^, and our observations might inform how proliferation is achieved in these conditions. Furthermore, the Gcn4p ortholog in mammals, Atf4, play important roles in cancer cell proliferation^66–68^, where many cancers continue to grow in apparently poor nutrient environments. Our study suggests how, in methionine rich (but otherwise amino acid limiting) conditions, Gcn4/Atf4 might function to promote growth, and not just help cells recover from nutrient stress. A separate, emerging question will be to understand how Gcn4p is itself regulated under these otherwise amino acid limited conditions, by methionine. Note that our studies would not have been possible without using prototrophic (“wild-type”) yeast strains to study responses to amino acids. Typically, studies utilize laboratory strains derived from an auxotrophic backgrounds (eg. S288C/BY4741), which require supplemented uracil, histidine, leucine and methionine for survival^69–74^, and where therefore overall amino acid homeostasis is severely altered. This precludes systematic experiments with amino acid limitation, such as those in this study.

We close by suggesting a possible metabolic cost based hypothesis for what might make methionine a strong growth cue. The *de novo* synthesis of methionine and its immediate metabolites (notably SAM) is exceptionally costly in terms of NADPH molecules invested^25,75–77^. Cells require at least 6 molecules of NADPH to reduce sulfur and synthesize a single molecule of methionine. Since biology has tied multiple anabolic processes to reductive biosynthesis (dependent on NADPH from the PPP), the availability of methionine might be an ancient signal to represent a metabolic state where reductive equivalents are sufficiently available for all other reductive biosynthetic processes as a whole.

## Materials and Methods

### Yeast strains and growth media

The prototrophic CEN.PK strain (referred to as wild-type or WT) was used in all experiments^78^. Strains with gene deletions or chromosomally tagged proteins (at the C-terminus) were generated as described. Strains used in this study are listed in Table S1.

The growth media used in this study are RM (1% yeast extract, 2% peptone and 2% lactate) and MM (0.17% yeast nitrogen base without amino acids, 0.5% ammonium sulfate and 2% lactate). All amino acids were supplemented at 2 mM. NonSAAs refers to the mixture of all standard amino acids (2 mM each) except methionine, cysteine and tyrosine.

The indicated strains were grown in RM with repeated dilutions (~36 hours), and the culture in the log phase (absorbance at 600 nm of ~1.2) was subsequently switched to MM, with or without addition of the indicated amino acids. For growth curves, the RM acclimatized cultures were used and diluted in a fresh medium with the starting absorbance of ~0.2 and the growth was monitored at the indicated time intervals.

### Western blot analysis

Approximately ten OD_600_ cells were collected from respective cultures, pelleted and flash-frozen in liquid nitrogen until further use. The cells were re-suspended in 400 μl of 10% trichloroacetic acid and lysed by bead-beating three times: 30 sec of beating and then 1 min of cooling on ice. The precipitates were collected by centrifugation, re-suspended in 400 μl of SDS-glycerol buffer (7.3% SDS, 29.1% glycerol and 83.3 mM Tris base) and heated at 100°C for 10 min. The supernatant after centrifugation was treated as the crude extract. Protein concentrations from extracts were estimated using bicinchoninic acid assay (Thermo Scientific). Equal amounts of samples were resolved on 4 to 12% Bis-Tris gels (Invitrogen). Coomassie blue–stained gels were used as loading controls. Western blots were developed using the antibodies against the respective tags. We used the following primary antibodies: monoclonal FLAG M2 (Sigma), and HA (12CA5, Roche). Horseradish peroxidase–conjugated secondary antibodies (mouse and rabbit) were obtained from Sigma. For Western blotting, standard enhanced chemiluminescence reagents (GE Healthcare) were used. ImageJ was used for quantification.

### Immunofluorescence measurements

Yeast cells were fixed with 3.7% formaldehyde, washed and resuspended in spheroplasting buffer (40 mM potassium phosphate buffer, pH 6.5; 0.5 mM MgCl_2_; 1.2 M sorbitol). Spheroplasts were prepared by zymolyase (MP Biomedicals, 08320921) treatment and spread on a slide pretreated with 50 μl of 1 mg/ml polylysine (Sigma-Aldrich, P6407). Gcn4-HA was stained with the mouse monoclonal anti-HA (12CA5) primary antibody (Roche, 11583816001) and Alexa Fluor 488-conjugated Goat anti-Mouse IgG (H+L) secondary antibody (Thermofisher, A32723). DNA was stained with 1µg/ml DAPI for 2 minutes, washed and mounted in Fluoromount-G (Southern Biotech, 0100–01). The cells were imaged using Olympus FV1000 confocal microscope.

### RNA-seq analysis

Total RNA from yeast cells was extracted using hot acid phenol method ^79^. The quality of RNA was checked on Bioanalyzer using an RNA 6000 Nano kit (Agilent) and the libraries were prepared using TruSeq RNA library preparation kit V2 (Illumina). The samples were sequenced on Illumina platform HiSeq2500. The raw data is available with NCBI-SRA under the accession number SRP101768. Genome and the annotation files of *S. cerevisiae* S288C strain were downloaded from Saccharomyces Genome Database (SGD; http://www.yeastgenome.org/). 100-mer, single-end reads obtained from RNA sequencing experiments were mapped to the S288C genome using Burrows Wheeler Aligner (BWA) ^80^. Mapped reads with the mapping quality of ≥ 20 were used for the further analysis. The number of reads mapped to each gene was calculated and the read count matrix was generated. The read count matrix was fed into EdgeR, a Bioconductor package used for analyzing differential gene expression ^81^. Genes which are differentially expressed by at least 3-fold with the p-value of <0.0001 were considered for further analysis. Normalized gene expression was calculated by dividing the number of reads by the gene length and the total number of reads for those samples, then dividing each of these values with the mode of its distribution ^82^. Absolute expression levels of the genes between the replicates are well correlated with the Pearson correlation coefficient (R) values more than 0.99 (see Figure S2). Mapping of genes to the related pathways and gene ontology analysis were carried out using public databases such as Yeastcyc ^83^, GeneCodis ^84–86^ and SGD ^87^.

### Metabolite extractions and measurements by LC-MS/MS

Cells were grown in RM for ~36 hours and transferred to MM with and without methionine for the indicated time. After incubation, cells were rapidly harvested and metabolite extracted as described earlier ^21^. Metabolites were measured using LC-MS/MS method as described earlier ^38^. Standards were used for developing multiple reaction monitoring (MRM) methods on Thermo Scientific TSQ Vantage triple stage quadrupole mass spectrometer or Sciex QTRAP 6500. For positive polarity mode, metabolites were separated using a Synergi 4μ Fusion-RP 80A column (150 × 4.6 mm, Phenomenex) on Agilent’s 1290 infinity series UHPLC system coupled to mass spectrometer. Buffers used for separation were: buffer A: 99.9% H_2_O/0.1% formic acid and buffer B: 99.9% methanol/0.1% formic acid (Flow rate, 0.4 ml/min; T = 0 min, 0% B; T = 3 min, 5% B; T = 10 min, 60% B; T = 10.1 min, 80% B; T = 12 min, 80% B; T = 14 min, 5% B; T = 15 min, 0% B; T = 20 min, stop). For negative polarity mode, metabolites were separated using a Luna HILIC 200A column (150 × 4.6 mm, Phenomenex). Buffers used for separation were: buffer A: 5 mM ammonium formate in H_2_O and buffer B: 100% acetonitrile (flow rate: 0.4 ml/min; T = 0 min, 95% B; T = 1 min, 40% B; T = 7 min, 10% B; T = 11 min, 1% B; T = 13 min, 95% B; T = 17 min, stop). The area under each peak was calculated using Thermo Xcalibur software (Qual and Quan browsers).

### ^15^N- and ^13^C- based metabolite labelling experiments

For detecting ^15^N-label incorporation in amino acids and nucleotides, ^15^N-ammonium sulfate with all nitrogens labelled (Sigma- Aldrich) was used. For ^13^C-labeling experiment, ^13^C- lactate with all carbons labelled (Cambridge Isotope Laboratories) was used. In the labelling experiments, 0.5X refers to 0.25% ammonium sulfate or 1% lactate. All the parent/product masses measured are enlisted in Table S2. Amino acid measurements were done in the positive polarity mode. For all the nucleotide measurements, release of the nitrogen base was monitored in positive polarity mode. For the ^13^C-label experiment, the phosphate release was monitored in negative polarity mode. Under these conditions, the nitrogen base release cannot be monitored here as the nitrogen base itself has carbon skeleton, which will complicate the analysis. The HPLC and MS/MS protocol was similar to those explained above.

### GDH assays

Glutamate dehydrogenase activity was measured as described in ^88^, with some modifications. Yeast cells were lysed by bead-beating in lysis buffer (100 mM potassium phosphate buffer, pH 7; 5% glycerol; 1 mM PMSF; 0.1% Tween-20; 1 mM EDTA; 1 mM 2-mercaptoethanol). NADP-dependent activity was measured by monitoring oxidation of NADPH (assay buffer: 100 mM Tris-HCl, pH 7.2; 10 mM 2-ketoglutarate, pH adjusted to 7.2; 100 mM ammonium chloride; 0.1 mM NADPH) at 340 nm. Protein concentrations from extracts were estimated using bicinchoninic acid assay (Thermo Scientific). One enzyme unit corresponds to the amount of enzyme required to oxidize one μmol of NADPH min^-1^ at room temperature.

### Statistical analysis

In most experiments, Student’s t-test was applied for calculating the p-values. Wherever necessary, other tests were applied and indicated accordingly.

## Acknowledgements

We acknowledge Dhananjay Shinde, Padma Ramakrishnan and the NCBS/inStem/C-CAMP Mass spectrometry facility for LC-MS/MS support, Avadheesh Pandit and the NCBS/inStem/C-CAMP next-generation sequencing facility for assistance with library preparation. We thank Utpal Banerjee, Mark Sharpley, Marco Foiani, Christopher Bruhn, Krishnamurthy Natarajan, Arati Ramesh and PJ Bhat for critical comments on this manuscript. This work was supported by a Wellcome Trust-DBT IA intermediate fellowship (IA/I/14/2/501523) and inStem/DBT institutional support to SL. ASW, RG, RS acknowledge bridging fellowships (from inStem), and ASW, RG and RS are supported by DST SERB- national postdoctoral fellowships (PDF/2015/000225, PDF/2016/000416 and PDF/2016/001877 respectively).

